# A narrow ratio of nucleic acid to SARS-CoV-2 N-protein enables phase separation

**DOI:** 10.1101/2024.04.10.588883

**Authors:** Patrick M. Laughlin, Kimberly Young, Giovanni Gonzalez-Gutierrez, Joseph C.Y. Wang, Adam Zlotnick

## Abstract

SARS-CoV-2 Nucleocapsid protein (**N**) is a viral structural protein that packages the 30kb genomic RNA inside virions and forms condensates within infected cells through liquid-liquid phase separation (**LLPS**). N, in both soluble and condensed forms, has accessory roles in the viral life cycle including genome replication and immunosuppression. The ability to perform these tasks depends on phase separation and its reversibility. The conditions that stabilize and destabilize N condensates and the role of N-N interactions are poorly understood. We have investigated LLPS formation and dissolution in a minimalist system comprised of N protein and an ssDNA oligomer just long enough to support assembly. The short oligo allows us to focus on the role of N-N interaction. We have developed a sensitive FRET assay to interrogate LLPS assembly reactions from the perspective of the oligonucleotide. We find that N alone can form oligomers but that oligonucleotide enables their assembly into a three-dimensional phase. At a ∼1:1 ratio of N to oligonucleotide LLPS formation is maximal. We find that a modest excess of N or of nucleic acid causes the LLPS to break down catastrophically. Under the conditions examined here assembly has a critical concentration of about 1 µM. The responsiveness of N condensates to their environment may have biological consequences. A better understanding of how nucleic acid modulates N-N association will shed light on condensate activity and could inform antiviral strategies targeting LLPS.

## Introduction

SARS-CoV-2 has caused 771 million infections and 7 million deaths since its emergence in 2019.(1) Highly effective vaccines have ameliorated the morbidity and mortality of this disease. However, the continued evolution of SARS-CoV-2 within humans and the wide array of documented animal reservoirs threatens a return to pandemic conditions.(2) Effective antivirals (e.g. Paxlovid) are available. However, Paxlovid requires early administration(3) and is cross-reactive with several common medications.(4) Furthermore, existing medications are ineffective for treating long COVID, which is associated with persistent reservoirs of SARS-CoV-2.(5)

N-protein (N) is the most abundant viral protein during a typical SARS-CoV-2 infection and is critical to nearly every step of the viral life cycle.(6) In the core of the mature virion, N forms 15 nm, roughly sperical assemblies with the viral genome that imply conserved architectures and protein-protein interactions.(7–9) After entering a host cell, N complexes must dissociate from the viral genome to enable primary translation. Within viral replication factories, N enables replication of the full-length genome through interactions with the viral replicase.(10, 11) Genomic RNA (∼30 kb) must be segregated from contaminating host and viral RNA’s in the cytoplasm, an ill-defined process that depends on liquid-liquid phase separation (LLPS) of N.(12, 13) Phase-separated condensates provide a scaffold for budding of new virions through interactions of N with the viral Matrix protein.(14, 15) N also has several immunomodulatory roles. It inhibits multiple pattern recognition receptors to downregulate cytokine production.(16–18) Paradoxically, N stabilizes inflammasomes and disassembles P-bodies to stimulate a pro-inflammatory response.(19, 20) LLPS is probably critical to many of these functions. N is displayed on the surface of the host cell, where it sequesters several chemokines.(21) Commensurate with these accessory roles, impaired clearance of N is associated with worse patient outcomes.(22, 23)

N is comprised of two ordered domains flanked by three intrinsically disordered domains (IDR’s; Figure 1A). The C-terminal ordered domain (CTD) forms a dimerization interface and also binds nucleic acid.(24) The N-terminal ordered domain (NTD) binds RNA specifically and non-specifically.(25, 26) The IDR linking the NTD and CTD (LKR) incorporates a leucine-rich helix that appears to participate in protein-protein interactions and a serine/arginine-rich region that can be phosphorylated to modulate LLPS formation.(11, 24, 27–29)

**Figure 1.**
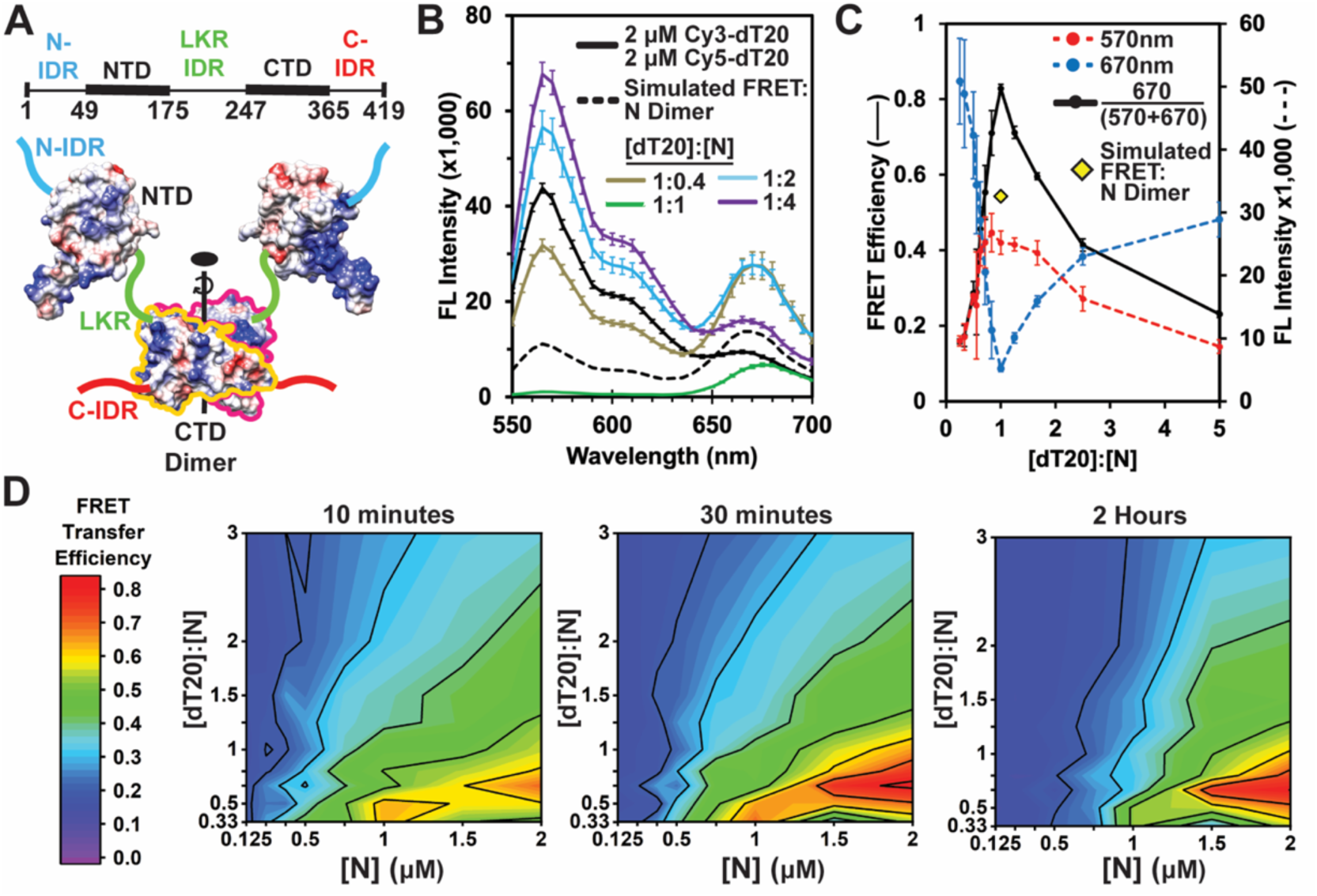
N packages Cy3- and Cy5-labeled nucleic acids in a multiprotein complex. (A) Schematic of an N dimer, with coulombic surfaces of NTD (PDB 8IQJ) and CTD (PDB 6WZO). (B) Emission spectra (excitation at 520nm) of labeled dT20 (2 µM each of Cy3- and Cy5-dT20) with different concentrations of N. Values in the legend represent the molar ratio the total dT20:N. Note that at 4 µM N (1:1) we observe almost complete loss of donor emission and substantial acceptor fluorescence (green line). This observation is distinctly different from a prediction for this condition, based on a single N dimer with two bound oligomers (dashed line). This implies formation of a larger complex. (C) The emission from donor (blue line) and acceptor (red line) shows a minimum near a molar ratio of 1 dT20 to 1 N. This minimum corresponds to a maximum in FRET efficiency (green line). The yellow diamond corresponds to the FRET efficiency predicted for an N dimer bound to two dT20s and not associated with any other N dimers. (D) FRET efficiency evolves over time as seen in phase diagrams of the [dT20]:[N] ratio versus [N] measured at 10 minutes, 30 minutes, and 2 hours. Phase diagrams also show evidence of a critical concentration near 0.5 µM N for formation of the high FRET efficiency dT20:N complex.

Proteins with IDR’s tend to undergo LLPS because they can interact with each other or with substrates such as RNA in a fluid manner.(30) These multivalent interactions lead to oligomerization and phase separation from bulk solution into membraneless organelles, whose formation must be reversible and responsive to their environment.(31) Dissecting the structural basis of LLPS is notoriously difficult due to its inherent aperiodicity and structural heterogeneity. For N, this is further compounded by its strong interactions with nucleic acid. While LLPS could be mediated entirely through nucleic acid, optical tweezer experiments with N show multi-step binding to DNA followed by “compaction”, which supports the necessity of protein-protein interactions.(32) However, the relative contribution of protein-protein versus protein-nucleic acid interations to N LLPS is unknown.

In this study, we examine the behavior of N LLPS *in vitro*. We show that N forms complexes with an oligonucleotide that phase separate at narrow ratios of oligonucleotide to N. Phase-separated condensates are dynamic and can grow over time. They are responsive to their environment and can be disrupted by addition of excess N or oligonucleotide. Finally, we demonstrate the importance of protein-protein interactions to N multimerization, which occur with or without oligonucleotide. The presence of oligonucleotide facilitates linkages between protein complexes that drive LLPS.

## Results

### A short nucleic acid promotes N self-association

Our broad focus is to understand the contribution of protein-protein interactions to phase separation of N with nucleic acid. We first investigated the range of conditions where N co-assembles with nucleic acids. For these studies we use ssDNA oligonucleotides with the sequence of twenty consecutive Thymidines (dT20). A 20-nucleotide DNA oligomer was previously shown to be the minimum length for inducing N self-assembly.(33) The short length of dT20 emphasizes the effect of protein-protein interactions to self-association and minimizes the contribution of cooperativity arising from a longer nucleic acid lattice.(34) Thymidine was used as it has negligible base stacking and does not form secondary structures.(35) For fluorescence experiments, dT20 was 5’ end-labeled with Cy3 or Cy5 (Cy3-dT20 or Cy5-dT20, respectively). The spectral properties of the Cy3 and Cy5 labels allow for direct measurement of nucleic acid binding and packaging by N.

Care was taken that the N used in these studies has little or no contaminating nucleic acid from the expression system. The purification includes a PEI precipitation to remove nucleic acid. The resulting protein has an A260/A280 absorbance ratio of 0.52, consistent with an absence of detectable nucleic acid.(36) Though N is usually thought of as a dimer with two major nucleic acid binding sites, for clarity in discussing molar ratios of N and oligonucleotide, in this paper we define N concentration in terms of monomer.(33)

To examine formation of N complexes driven by interaction with short oligonucleotides, we leveraged Förster Resonance Energy Transfer (FRET). We prepared samples with different concentrations of N and a constant concentration of the two labeled dT20s (1:1 Cy3-dT20:Cy5-dT20) (Figure 1B). We reasoned that formation of a complex with two or more dT20s would bring Cy3 and Cy5 within their Förster distance (R_0_=5.3 nm) and cause FRET. When there is no N present, the emission spectrum observed with excitation at 520 nm (the excitation maximum for Cy3) shows no detectable FRET. With a dT20:N molar ratio of 1:0.4, we observe significant FRET, shown by a loss of Cy3 donor emission at 570 nm and an increase in acceptor emission for Cy5-dT20 at 670nm. Strikingly, at a 1:1 ratio of dT20 to N there is a near-total elimination of Cy3 fluorescence with a substantial amount of Cy5 emission. Higher concentrations of N result in a recovery of Cy3 fluorescence while Cy5 fluorescence had an initial increase followed by a gradual decrease. These results can be summarized by focusing on the peak emission wavelengths or on FRET efficiency (eqn 1, methods; Figure 1C). We note that the dT20:N ratio that yields the highest FRET efficiency changes depending on the dT20 concentration (Supplementary Figure 1A). Further, the behavior of dT20 is indistinguishable from an rU20 ssRNA (Supplementary Figure 2).

The rise and fall in FRET efficiency suggests a phase transition requiring a narrow dT20:N ratio. In particular, we focused on the spectrum of the 1:1 complex where donor emission was suppressed (Figure 1B-C) and compare this to predictions based on three models. For these predictions, we factored in Cy3-Cy3 and Cy5-Cy5 self-quenching and protein-induced fluorescence enhancement with control experiments where Cy3-dT20 or Cy5-dT20 mixed with unlabeled dT20 was combined with N (Supplemental Figure 1B). As a boundary condition, we assume that FRET transfer is 100% efficient for a complex with both Cy3 and Cy5 oligos (i.e. acceptor emission and no donor emission). In the simplest scenario we consider dT20 binding to an N dimer (Supplementary Figure 3A). Here we expect 25% of N dimers to be bound by two Cy3-dT20s, 25% to be bound by two Cy5-dT20s, and 50% to be bound to one of each labeled DNA oligo. The expected fluorescence spectrum is qualitatively and quantitatively different from our observed 1:1 spectrum (Figure 1B; compare black dashed line and green line). The calculated spectrum has significant donor emission from dimers with two Cy3-dT20s and less Cy5 emission due to the fact only 50% of N dimers are bound to a FRET pair. The much higher donor emission and lower FRET efficiency of our simulation (Figure 1C; yellow diamond with black border) relative to our experimental observations suggest a larger complex.

Given that a single dimer is inadequate to explain our results, we considered an N tetramer and a higher-order model. We first consider two N dimers (a tetramer) bound to four dT20s (Supplementary Figure 3B). In this model, 87.75% of tetramers would include both Cy3- and Cy5-dT20 and generate some FRET. 6.25% of tetramers would have four C5-dT20s. Maximal Cy3 emission (per complex) would arise from the 6.25% of tetramers bound to four Cy3-dT20s, which would yield far more Cy3 emission than observed. A higher order model comprised of many dimers (or tetramers) inherently resembles an LLPS (Supplementary Figure 3C). Phase separated condensates would contain a mixture of both oligos and thus yield high FRET because very few Cy3-dT20s would escape proximity to one or more Cy5 acceptors. While LLPS explains our observations (discussed later), the FRET assay cannot differentiate the size of LLPS condensates.

To gain a better understanding of the range of conditions for maximal FRET and to examine its N concentration dependence, we constructed phase diagrams of dT20:N versus [N] (Figure 1D). We observed the highest FRET efficiency at a 1:1.5 ratio of dT20:N. This maximum differs from the 1:1 ratio in the one dimensional version of this experiment (Fig 1B) suggesting either that these experimental conditions do not support tight binding and/or that association can lead to kinetic traps that impede equilibration. In support of the assertion that our concentrations are close to the K_D_ of N for dT20, we saw that a 4 μM concentration of dT20 yielded a maximum signal at a 1:1 ratio with N, while lower concentrations of dT20 required a molar excess of N to yield the same signal (Supplementary Figure 1A). We also observe that FRET efficiency increases over time (Figure 1D), suggesting continued growth of large N complexes. FRET fell off steeply on either side of the maximal signal, though not to baseline.

This steep falloff suggests a lack of large oligomers (where energy transfer is highly efficient) and a predominance of smaller complexes. These hypotheses are tested in later experiments. At a constant dT20:N ratio, we observed that maximal FRET required ≥1.25 µM N protein (Figure 1D). Formation of large complexes, a crystal or an LLPS, should be characterized by a critical concentration.(37–39) In a phase where subunits can freely equilibrate with bulk solvent, all subunit above the critical concentration is bound in the separated phase. Critical concentration is a function of the subunit-subunit interaction energy and the multivalency of the sububnit. The appearance of FRET signal at higher [N] is consistent with a critical concentration of LLPS assembly that is modulated by the dT20 concentration.

### dT20 packaging occurs in conjunction with liquid-liquid phase separation

Due to the length of our dT20 (135 Å) and the Förster distance of our Cy3-Cy5 FRET pair (R_0_ = 53 Å), our FRET experiments are focused on local interactions. To gain a more global understanding of how dT20 drives N self-assembly, we examined N-dT20 complexes using light scattering and light microscopy.

The change in light scattering was visually obvious and responsive to the ratio of dT20 to N (Figure 2A). With N alone and up to a 1:1 molar ratio, samples in Eppendorff tubes were clear. At a 1.5:1 and 2:1 ratio, the samples were very cloudy. Higher ratios of dT20:N again produced clear samples. More quantitatively, turbidity at 320 nm followed the same trend (Figure 2B).

**Figure 2.**
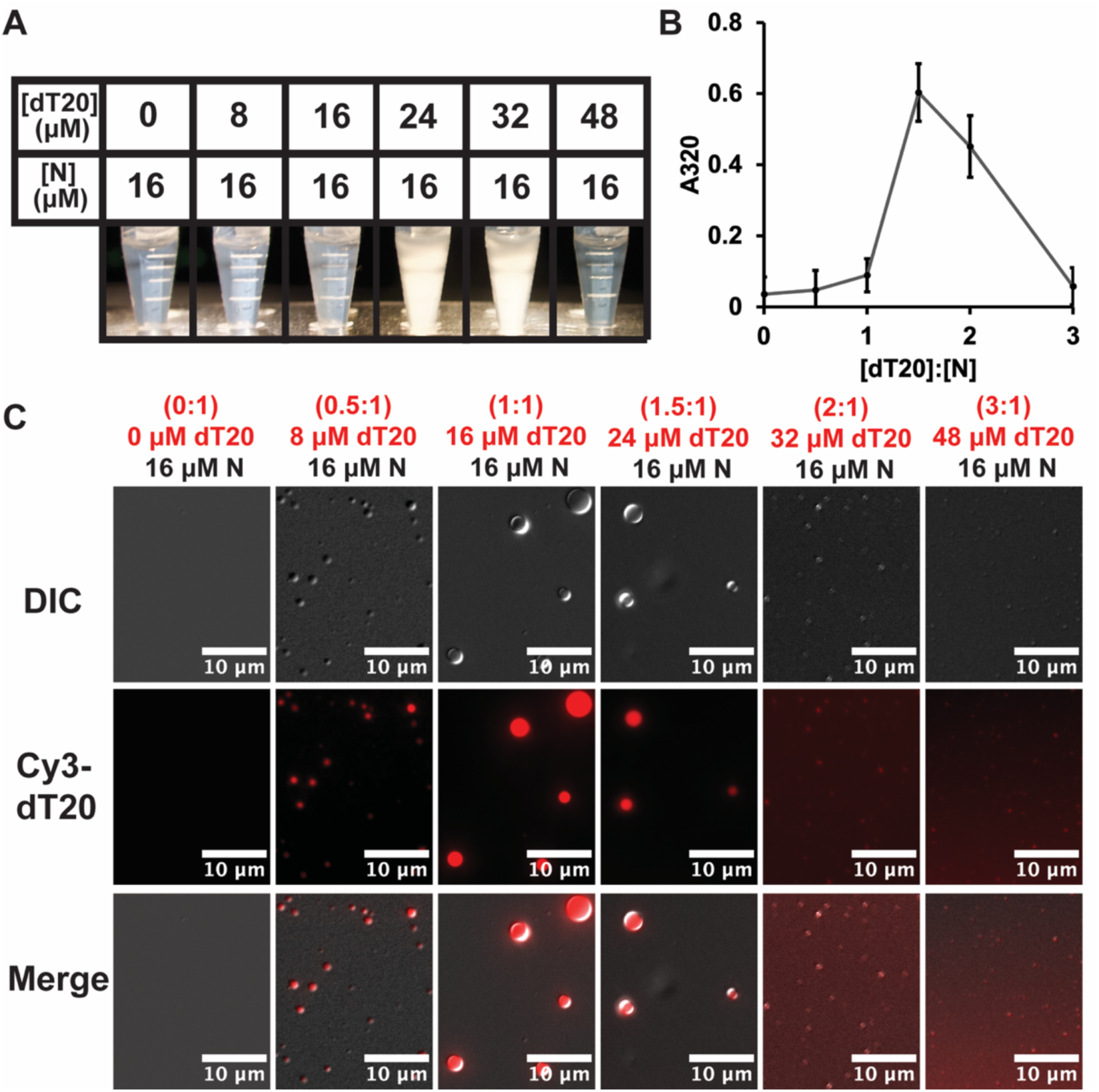
Liquid-Liquid Phase Separation occurs at a dT20:N ratio near 1:1. (A) Visual observation of cloudiness in Eppendorf tubes and (B) 320 nm turbidity measurements 15 minutes after mixing each sample indicate that a maximum in light scattering occurs at distinct ratios of dT20 to N. (C) LLPS was detected using differential interference contrast microscopy. Fluorescence of Cy3-dT20 showed localization to condensates. At 8 µM Cy3-dT20, few condensates formed. At a dT20:N ratio of 1:1 (16 µM dT20), many large condensates were observed. At higher concentrations of dT20, condensates were smaller while a background of Cy3 fluorescence suggests that some dT20 is not bound by N.

The complexes that scatter light were further characterized by differential interference contrast (DIC) microscopy, which revealed that elevated sample turbidity was due to liquid-liquid phase separations (Figure 2C). Without any dT20, we did not see any condensates. At a 0.5 dT20:N ratio, a few condensates were found. At 1:1 and 1.5:1 ratio we saw larger condensates that were also abundant. While there were still abundant condensates at a 2-fold excess of dT20, they were much smaller in size. We could not detect any LLPS at higher dT20 concentrations.

To verify that LLPS was due to nucleic acid-driven assembly, we correlated DIC microscopy with epifluorescence imaging of condensates formed by N and Cy3-dT20. We observed colocalization of fluorescence with condensate. At samples where Cy3-dT20 was in excess of the 1:1 ratio, we noticed substantial background signal for Cy3-dT20, indicating that excess oligo is not associated with the condensates.

Like the FRET assays (Figure 1), there is a sharp rise and fall in turbidity and condensate formation as the ratio of dT20 to N is changed. We observed maximum FRET efficiency at a dT20:N ratio of 1:2 at low dT20 concentrations and at 1:1 ratio at higher dT20 concentrations (Supplementary Figure 1A). In our turbidity assay, we see that a molar excess of dT20 between 1.25- and 1.5-fold is required for maximum signal. We note that in our microscopy data, a large number of condensates are observed at these conditions, but they are smaller in general. This disconnect between assays may arise from the transition from Rayliegh to Mie scattering, discussed in the following paragraphs.

### Phase-separated Condensates are dynamic and coalesce over time

Given the similarities between the pattern for turbidity and LLPS formation observed by DIC, we generated phase diagrams for turbidity (Figure 3A) like those generated with the FRET-based assay (Figure 1D). At 10 minutes, we see the same trend as in Figure 2B for maximum turbidity at a dT20:N of 1.25:1 and 1.5:1. Higher N concentrations led to higher turbidity at the same dT20:N ratios. We detected elevated turbidity at N concentrations as low as 1 μM. At higher concentrations of N, a broader range of dT20:N was permissive for LLPS. No changing turbidity was detected in samples with N alone.

**Figure 3.**
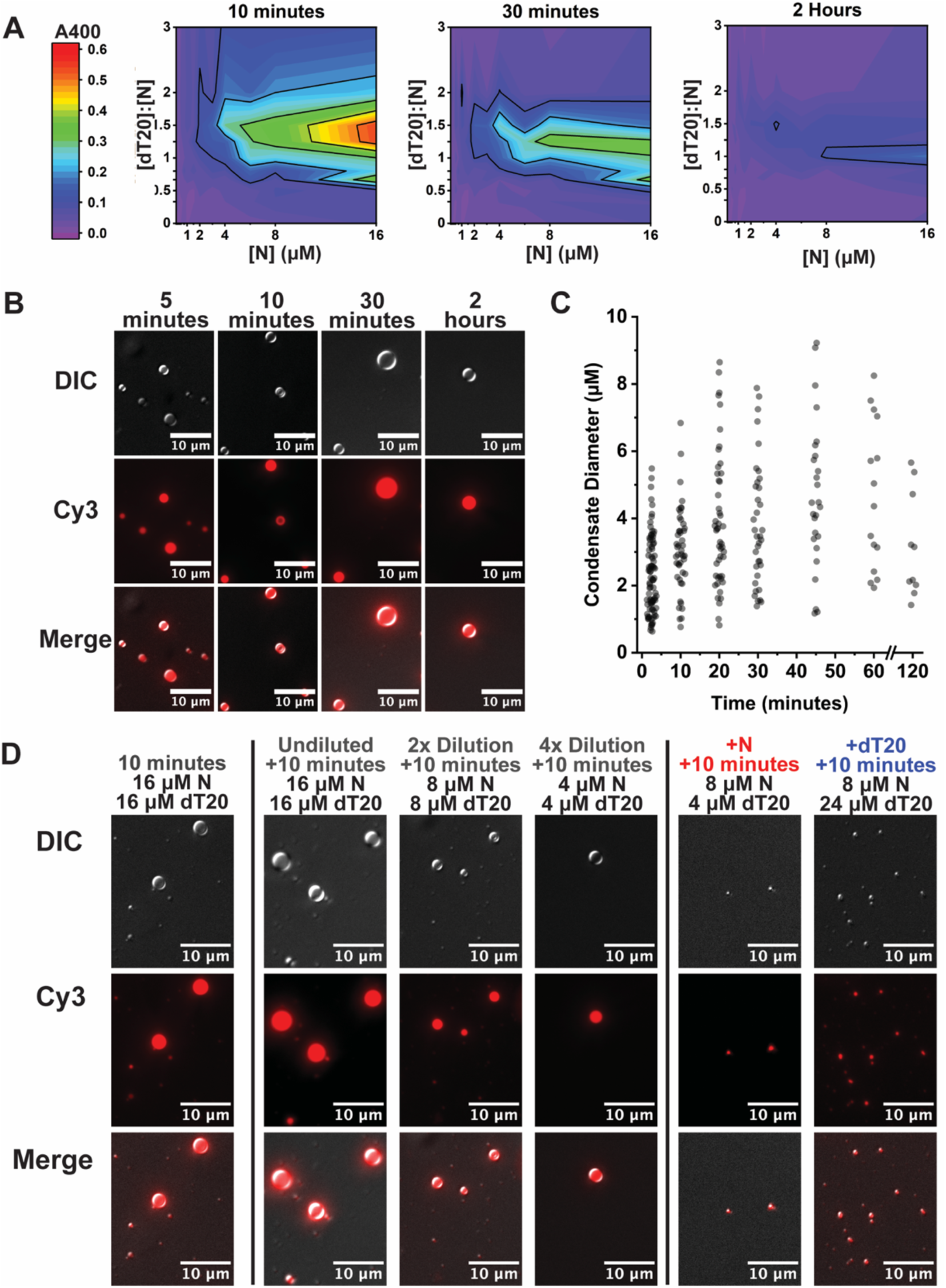
LLPS is concentration-dependent and dynamic. (A) 400 nm turbidity measurements indicate that a maximum in light scattering occurs at a 1.5:1 excess of dT20:N at 10 minutes. Turbidity falls sharply by 30 minutes. At 2 hours, turbidity falls further with a maximum at equimolar dT20:N. (B) Condensates were assembled at 16 µM Cy3-dT20, 16 µM N and visualized with differential interference contrast. Cy3-dT20 was localized to condensates, which grew over time. (C) Condensate sizes from (B) were determined by thresholding of Cy3 fluorescence intensity. Condensates grow until plateauing near 4 microns. (D) Condensates diluted with excess Cy3-dT20 or N are smaller than dilution with buffer alone and indicates destabilization caused by excess free subunit.

Counterintuitively, turbidity of the samples decreased very quickly. By 30 minutes, the signal had decreased by over 50% from the first measurement at 10 minutes for most conditions. By 2 hours the turbidity signal shifted to a maximum at 1:1 dT20:N. This is identical to the ratio that yielded maximum FRET in prior results (Figure 1B, C; Supplementary Figure 1A).

The decrease in turbidity may be attributable to a change in the scattering regimen of LLPS condensates. At early timepoints in assembly, where condensates are expected to be small, we suggest that samples are in the Rayleigh Scattering regimen, where turbidity is proportional to the weight-average molecular weight of the assembled products. As condensate diameter grows large compared to the wavelength of scattered light (>>5% the wavelength of incident light, >>20 nm), they enter the Mie Scattering regimen, which causes a decrease in light scattering. We tested the prediction of LLPS growth with DIC and epifluorescence (Figure 3B, C). At 10 minutes, many small condensates (<1 micron diameter) were observed that colocalized with Cy3-dT20. With longer incubation, we observed larger condensates that were fewer in number. We analyzed particle size and polydispersity through direct measurement and saw that condensates indeed grew over time until approaching a median diameter of ∼4 microns after 2 hours. At these sizes, Rayliegh theory is inadequate; Mie scattering thus accounted for the decrease in the observed scattering over time, as smaller condensates decrease in number due to growth or incorporation into larger preformed condensates. Efforts to visualize aggregates by negative stain and cryo electron microscopy were unsuccessful, probably because they were disordered, irregular, and large Because LLPS size is dynamic, we predicted that preformed condensates would be responsive to changes in their environment. In particular, because LLPS formation is very sensitive the the dT20:N ratio, we hypothesized that additions of free N or free dT20 would perturb preformed LLPS condensates more than dilution with buffer alone. To test this hypothesis, we first assembled condensates at an equimolar ratio of N to Cy3-dT20. After a preincubation, we diluted this stock with free N, free Cy3-dT20, or buffer. Despite dilution with buffer, preformed condensates were resistent to disassembly. However, addition of free Cy3-dT20 or N resulted in condensates that were much smaller than our control dilutions. Based on DIC microscopy, we see that dilution has little effect on condensate size (Figure 3D). The ability of excess N or dT20 to disrupt condensates could be of biological significance.

FRET, turbidity, and DIC microscopy illustrate that N will phase separate over a narrow range of ratios of dT20 to N when in excess of a critical concentration. Consistent with their dynamic nature, phase separations are disrupted by addition of excess N or dT20.

### N assembles through protein-protein as well as protein-oligonucleotide interactions

In these experiments, N does not appear to form a separate phase without dT20. This raises the question of whether protein-protein interactions are involved in LLPS. We turned to crosslinking to test for direct interaction between N-proteins.

We reasoned that salt bridges would be likely in oligomerization of N which is extremely polar and enriched in intrinsically disordered regions. 1-Ethyl-3-(3-dimethylaminopropyl) carbodiimide (EDC) forms a zero-length covalent linkage between a primary or secondary amine and a carboxylate (e.g. lysine and aspartate or glutamate). We add sulfoNHS to EDC reactions to form a more stable reaction intermediate and thus increase reaction efficiency. Similarly, for N to form a phase with nucleic acids, basic regions were likely to be in juxtaposition bound to a bridging nucleic acid. Thus, a second complementary crosslinker is bis(sulfosuccinimidyl)suberate (BS3), which links two primary amines with an 11.4 Å linker. Crosslinking was evaluated by SDS-PAGE (Figure 4A-D).

**Figure 4.**
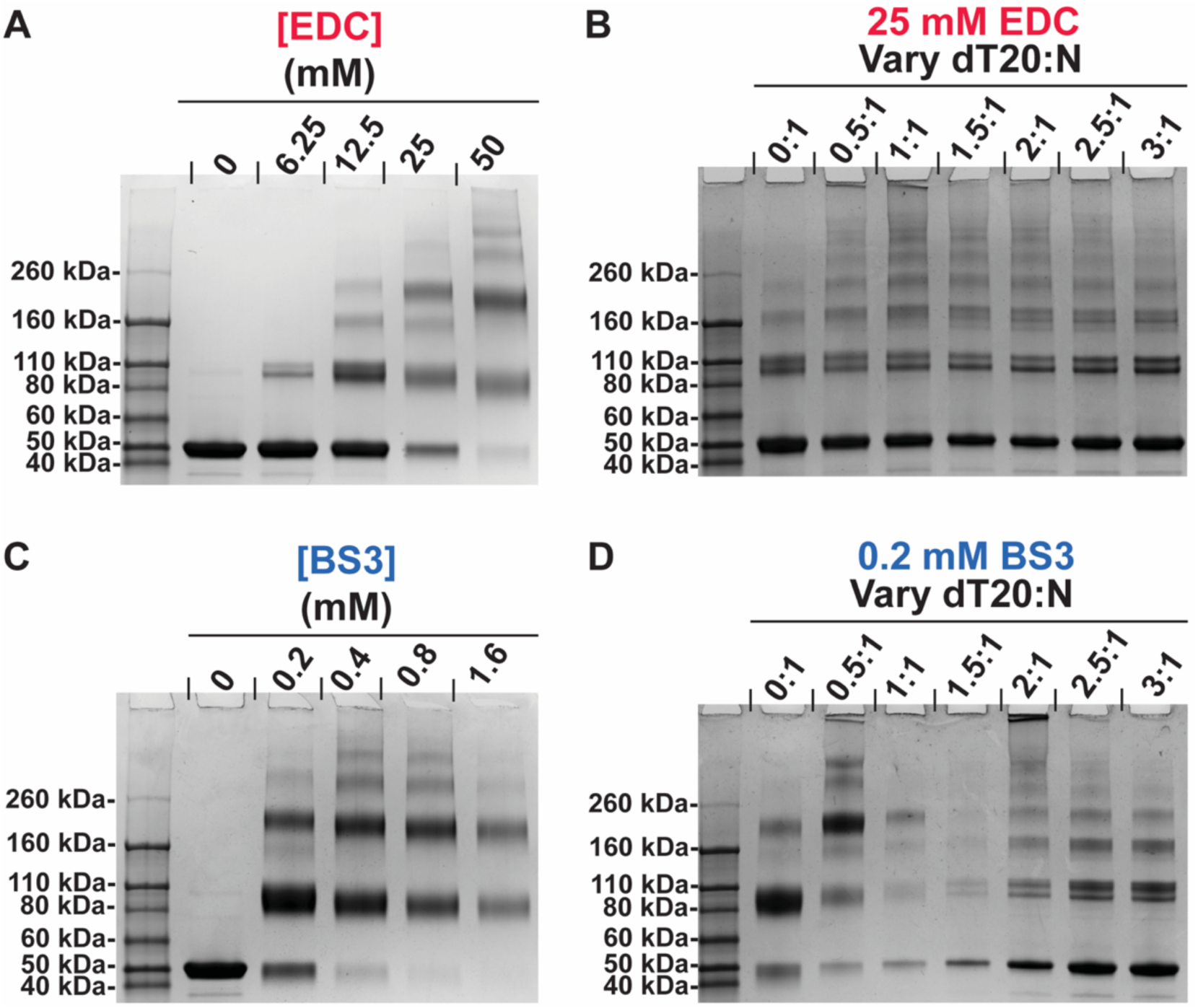
N constitutively assembles and is modulated by dT20. (A) EDC and (C) BS3 Crosslinking of 16 µM N without dT20 shows N self-assembly. Bands corresponding to monomer, dimer, tetramer, and higher-MW bands via SDS-PAGE. Crosslinking of 16 µM N in the presence of dT20 by (B) 25mM EDC and (D) 0.2mM BS3. The EDC treated sample shows higher MW bands that are more dominant in samples that phase separate. For BS3 crosslinked LLPS most of the material does not enter the gel, hence the weaker bands. For EDC crosslinking, sulfoNHS was added to 0.2x the concentration of EDC.

Both EDC and BS3 led to extensive crosslinking of N without any dT20 present. We observed a concentration dependent formation of dimer, trimer, tetramer, and higher-MW species (Figure 4A, 4C). To test for formation of spurious crosslinks, reactions were also run in the presence of 50 mM Tris, which should effectively compete with non-specific partners. With both crosslinkers we were able to stabilize a similar catalog of N oligomers even in the presence of Tris (Supplementary Figure 4A-B).

We investigated crosslinking in the presence of dT20. Samples from phase separations were largely associated with higher-MW bands. With EDC (Figure 4B), these samples led to a ladder of higher-MW bands that were not observed without dT20. BS3 (Figure 4D) was much more responsive to the concentration of dT20. Low concentrations of dT20 led to a novel ladder of intermediates. At dT20 concentrations consistent with LLPS formation, little N migrated into the gel consistent with it being trapped in a very large complex. The complexity arises in the presence of sufficient dT20 to disrupt an LLPS. Under these conditions, we observe a novel ladder of bands distinctly different from what was seen in the absence of dT20 or in low concentrations thereof.

These results show that N without nucleic acid can form a range of species. N self-association is frequent enough that crosslinkers can outcompete high concentrations of competitive inhibitor. When oligonucleotide is added, N undergoes further multimerization. Differences in the EDC and BS3 crosslinking pattern of phase separated condensates indicate that a unique set of protein-protein interactions are required for LLPS.

## Discussion

In this work, we focus on four observations. First, a narrow range of dT20:N enables LLPS. Second, condensates are disrupted by addition of excess N or dT20, which is suggestive of a biologically relevant responsiveness. This dynamic nature of LLPS may influence nucleic acid packaging, genome segregation and virus assembly, or release of free N for innate immune modulation. Third, phase separation requires a critical concentration of N. Though lacking any apparent structural order, LLPS growth is analogous to nucleated polymerization such as crystallization. Fourth, N assembles through both protein-protein and protein-nucleic acid interactions. However, neither set of interactions alone is sufficient for LLPS.

These behaviors inform a simple model in which protein-protein and protein-nucleic acid interactions control the N assembly landscape (Figure 5). With a 1:1 dT20:N ratio, we observed aggressive LLPS formation. This result is consistent with the observations of Zhao et al who observed that a 20-mer was long enough to induce assembly of N dimers and built a similar model.(33) All five domains of N contribute to nucleic acid interactions.(40, 41) We infer that each N monomer has two effective dT20 binding sites, or four sites per N dimer. As noted by Schuck and co-workers, a dT20 is sufficiently long to be bound by two N dimers.(33) At a 1:1 ratio, Each N dimer has two open and two filled sites and can participate in LLPS growth. In addition to nucleic acid crosslinks and dimerization by the ordered CTD, we observe additional protein-protein interaction by crosslinking, which are consistent with those identified from crystallographic and solution experiments.(14, 24, 27–29, 42, 43) Substantial published work indicates interactions via the leucine-rich region of the LKR IDR connecting the ordered NTD and CTD domains.(27, 28) Crystal contacts suggest CTD-CTD interaction.(44) Also the C-terminal IDR has been implicated a site of interaction.(24, 45) However, in the absence of nucleic acid these protein-protein interactions are only sufficient to support small complexes (Fig 4A,C).

**Figure 5.**
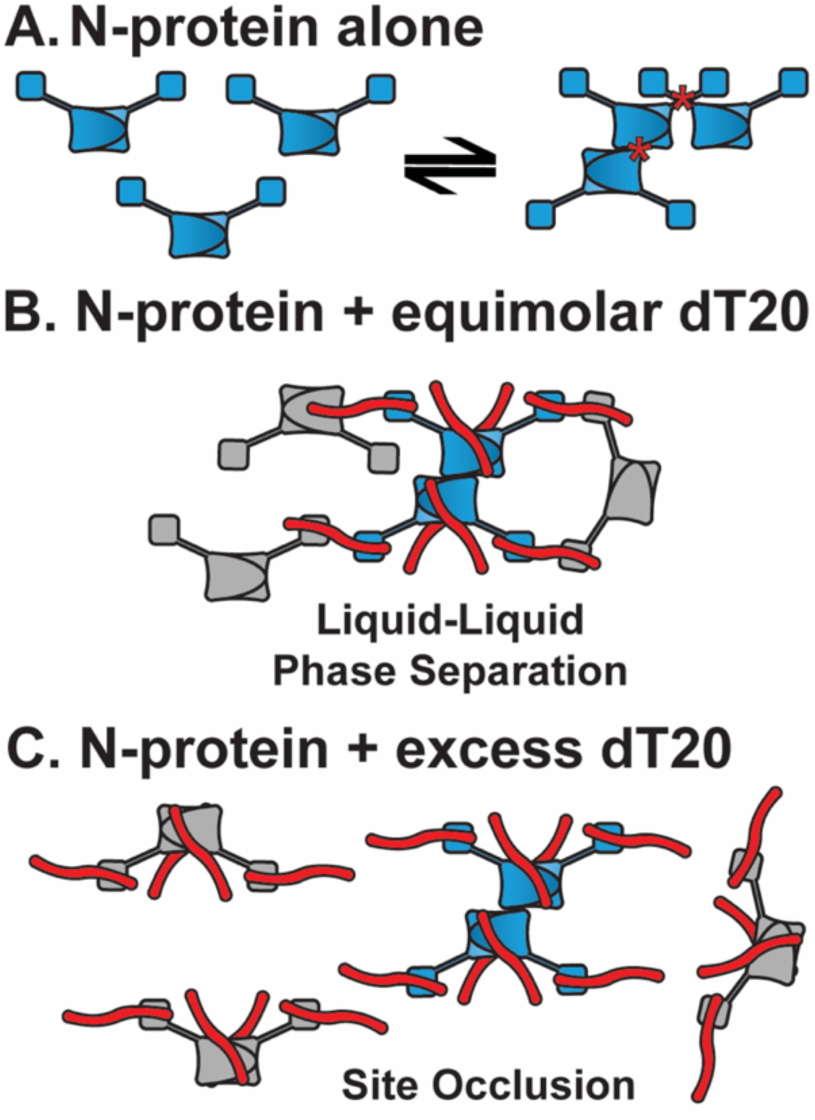
Incomplete saturation of N with nucleic acid is required for phase separation. (A) The cartoon of an N dimer based on the structure shown in Figure 1A; the N-IDR and C-IDR are omitted for clarity. N-protein self-association occurs stochastically without nucleic acid to form multimers that are LLPS-incompetent. In this diagram, oligomerization occurs through CTD-CTD interaction and mediated by L-rich helices located in the LKR (contacts denoted by the asterisks). (B) Each NTD has one oligonucleotide binding site and the CTD dimer has two sites. The dT20 oligonucleotide is long enough to form linkages between N dimers. With two dT20s per dimer, each dT20 can be shared by two dimers, enabling extensive network formation and phase separation. Shown is a central dimer of dimers with all nucleotide sites filled (blue) linked to partially filled surrounding dimers (gray). (C) Excess dT20 fills all sites and thus cannot accept an oligonucleotide from other N-protein dimers, disrupting LLPS connectivity. Even with nuleic acid sites filled, there can still be oligomerization by protein-protein interactions.

This model of an N dimer with four effective dT20 sites provides an explanation for LLPS growth and also for LLPS response to excess N and excess dT20. At the 1:1 ratio, where half of all sites are empty, the periphery of the LLPS will be comprised of open sites and “ends” of dT20 molecules. It will be maximally receptive to adding more 1:1 dimers and complexes of dimers. We observe continued growth under these conditions. Unlike crystals, which do not typically merge, an LLPS can adjust its surface geometry to encourage fusion. Barring a physical constraint, the larger the LLPS the larger the fraction of buried subunits and the lower the energy of the complex. Subunits on the interior may have incomplete or sub-optimal contacts, as with any liquid. The relative affinity of the two dT20 binding sites may influence the kinetics of LLPS. For free N dimers, bound dT20 is simply not shared with N in another dimer. In the context of LLPS, all sites are engaged due to “sharing” of dT20 between adjacent N proteins. Links that promote LLPS may be kinetically inhibited by dT20 repositioning despite favorable thermodynamics. In our turbidity phase diagrams (Figure 3A), we observed that the dT20:N ratio that yielded maximum turbidity changed from 1.5:1 at 10 minutes to 1:1 after two hours. The higher turbidity of the 1.5:1 ratio at shorter timescales may be explained by restoration of dT20 to the weaker (but LLPS-critical) binding site.

Addition of excess N (or dT20) to an LLPS is predicted to result in an LLPS coated with unsaturated or oversaturated N and lead to stripping off peripheral subunits. If N does not infiltrate the LLPS, loss of subunits will expected to proceed from the exterior. If N does infiltrate the complex, we can anticipate accumulation of defects in the LLPS lattice, a percolation threshold, and a sudden collapse of the LLPS. (46–48) The susceptibility of LLPS integrity to an excess of N may allow the LLPS to respond to changes in the biological milieu and progression through the viral replication cycle. Consider a situation where the relative amount of N in the host cell increases. As virus replication proceeds, the RNA:N ratio decreases changing LLPS stability and leading to virus assembly.

A characteristic of a polymer (e.g. a crystal) that is applicable to an LLPS is that, for a set of conditions, its assembly can be described in terms of a critical concentration. In the low ionic strength, short oligo conditions investigated in this study, the critical concentration is approximately 1 µM (Figure 1, 3). A critical concentration arises when subunits are free to equilibrate between association with free ends and bulk solution.(37) Thus, a critical concentration is a dissociation constant described as the ratio of the second order binding subunit to LLPS and the first order release of subunit:

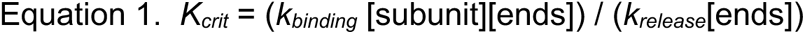

The value for *K_crit_* is a function of the microscopic association constants of interaction and the number of such interactions per subunit.(37, 49)

The contribution of N-N interactions to oligomerization provides thermodynamic support for assembly of amorphous LLPS. These interactions may also play roles in other structural functions of N. In tomographic studies of virions, N was found to form small 15nm diameter complexes, about 40 per virion, that are believed to compact and organize the viral genome, analogous to nucleosomes.(7–9) We used crosslinking to search for interaction in the absence of nucleic acid (Figure 4A,C) and found evidence of salt bridges (crosslinked by by EDC) and clusters of basic residues (crosslinked by BS3). To avoid serendipitous crosslinks, we added up to 50 mM Tris and nonthess saw essentially the same interactions. However, this self-assembly is limited, structurally or energetically, to a few subunits based on crosslinking and the absence of turbidity or LLPS (Figure 2A-B, Figure 3A). This lack of phase separation of free N protein, even in the presence of crosslinking agents, suggests that by itself N was not nucleating a three dimensional phase.(37) dT20 concentrations that induce N LLPS also stabilize N-dimerization and yield higher-mass crosslinks (Figure 4). Nonetheless, N multimerization in the absence of nucleic acid may provide avidity for nucleic acid engagement and further protein-protein interaction.

In our experiments, growth of LLPS shares many similarities to growth of a protein crystal. Both are comprised of repeating units that associate by weak contacts. In a protein crystal, interactions are persistent and each repeating cell has the same ensemble of contacts. In LLPS, the contacts are transient and a given subunit may only have a fraction of its possible contacts. The growth of a crystal or LLPS are governed by the strength of these contacts, which can be evaluated in terms of a critical concentration, the maximum concentration of free sununit at equilibrium. Growth of LLPS will continue until either the concentration of free subunit drops to the critical concentration or the surface is “poisoned” by defective subunits. In our FRET and turbidity assays, we observed high responsiveness to the N concentration. We also observe that “poisoning” of LLPS by excess N or dT20 can disrupt condensates much more severely than an equivalent dilution of buffer alone.

An N protein LLPS must involve both protein-protein and protein-nucleic acid interactions. While protein-protein interactions are sufficient to drive limited multimerization, LLPS could not be detected with N alone. While nucleic acid “bridging” interactions driven by exclusively N-dT20 interactions could theoretically drive LLPS, the growth pattern with dT20 (Figure 4C-D) showed many similarities to N alone. We suggest that the presence of identical multimerized species with and without dT20 implies a substantial role for protein protein interactions in the N LLPS pathway. The requirement for dT20 for LLPS suggests that longer range interactions between separate quasisimilar “unit cells” are sewn together by nucleic acid scaffolds. Taken together, these results suggest that modulation of N-N or N-nucleic acid contacts (i.e. by a small molecule) could alter the behavior of N condensates and disrupt multiple steps of the SARS-CoV-2 replication cycle.

## Methods

### Preparation of Material

DNA oligomers – dT20, Cy3-labeled dT20 (Cy3-dT20), and Cy5-labeled dT20(Cy5-dT20) – were purchased from Integrated DNA Technologies. For Cy3-dT20 and Cy5-dT20, the fluorophore was at the 5’ end. Concentrations were based on absorbance at 260 nm using extinction coefficients of 162,600 M^-1^cm^-1^ for unlabeled dT20, 167,500 M^-1^cm^-1^ for Cy3-dT20, and 172,600 M^-1^cm^-1^ for Cy5-dT20.

To create an N expression platform, we utilized the coding sequence for SARS-CoV-2 N (GenBank: NC_045512.2; Integrated DNA Technologies; ref 10006625), and cloned this sequence into a pET-24a(+) plasmid to introduce a C-terminal HisTag. After sequencing (Eurofins), the plasmid was transformed into BL21(DE3) cells (New England Biosciences) for protein expression. Cells were grown in Terrific Broth with 0.05 mg/mL kanamycin. Upon reaching an OD of 0.6-0.8, expression was induced with 1 mM IPTG for 18 hours at 23^°C^. Cells were cooled to 4^°C^ and pelleted by centrifugation (15 minutes at 10,000*xg*), then stored at −80^°C^.

The following protein purification steps were all performed at 4^°C^. Frozen cell pellets were resuspended in Lysis Buffer (50 mM HEPES [pH 7.5], 100 mM NaCl, 1 mM NaF, 0.1% βME with Roche EDTA-free protease inhibitor tablets; ref A32965) and lysed by Emulsification (Avestin). Cell debris was centrifuged at 10,000*xg* for 15 minutes. To the supernatant, polyethyleneimine (PEI; branched, MW 2000; Polysciences) was added to a final concentration of 1% w/v to precipitate nucleic acids, which were pelleted by centrifugation at 10,000*xg* for 10 minutes. Using ammonium sulfate (40% saturation), N was precipitated from the supernatant of the PEI precipitation and pelleted by centrifugation at 10,000*xg* for 10 minutes. The pellet was resuspended in NiNTA Binding Buffer (20 mM Tris [pH 7.5], 500 mM NaCl, 60 mM Imidazole, 0.1% βME) and loaded onto a 5 mL HisTrap HP column (Cytiva) equilibrated in the same buffer. The column was washed with 15 mL of Binding Buffer, followed by 15 mL with 5% NiNTA Elution Buffer (20 mM Tris [pH 7.5], 500 mM NaCl, 600 mM Imidazole, 0.1% βME). The protein was then eluted with 45% NiNTA Elution Buffer, which removed all the bound protein. Protein Fractions were diluted five-fold with Dilution Buffer (50 mM HEPES [pH 7.5], 0.1% βME) to lower the ionic strength before loading onto a 5 mL HiTrap SP FF column (Cytiva) pre-equilibrated with HiTrap Binding Buffer (50 mM HEPES [pH 7.5], 100 mM NaCl, 0.1% βME). The column was washed with 30 mL of HiTrap Binding Buffer, then 30 mL of 15% HiTrap Elution Buffer (50 mM HEPES [pH 7.5], 2 M NaCl, 0.1% βME). Protein was eluted with 30% HiTrap Elution Buffer and collected for dialysis into storage buffer (50 mM HEPES [pH 7.5], 150 mM NaCl, 0.1% βME). N stocks were highly pure and no nucleic acid contamination was present (A260/A280=0.53).

### Förster Resonance Energy Transfer (FRET) Assay

Samples for fluorescence measurements were prepared in black 384-well microplates (Greiner Bio-One; ref 784900) by a BioMek FX-P liquid handling robot (Beckman Coulter) at 23^°C^. Assays were performed on separate days for statistical analysis. Fluorescence data were collected using a Synergy H1 plate reader (BioTek) using excitation wavelengths of 520 nm or 600 nm. Using an excitation wavelength of 520 nm, the FRET transfer efficiency was defined as:

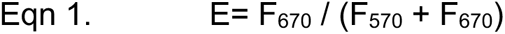

Wherein F is the fluorescence emission at the subscripted wavelength and E is FRET efficiency. Wells containing Cy3-dT20 and unlabeled-dT20 or Cy5-dT20 and unlabeled-dT20 were used quantify the effect of Cy3-Cy3 and Cy5-Cy5 self-FRET, as well as protein-induced fluorescence enhancement, to our signal.

For the experiment in Figure 1B-C and Supplementary Figure 1A-B, three oligonucleotide stocks (2 μM Cy3-dT20, 2 μM Cy5-dT20; 2 μM Cy3-dT20, 2 μM unlabeled dT20; 2 μM Cy5-dT20, 2 μM unlabeled dT20; each in 50 mM HEPES [pH 7.5], 0.1% βME) and one N stock (16 μM N in 50 mM HEPES [pH 7.5], 20 mM NaCl, 0.1% βME) were prepared. For the experiment in Figure 1D, different concentrations of oligonucleotide (3 μM Cy3-dT20, 3 μM Cy5-dT20; 3 μM Cy3-dT20, 3 μM unlabeled dT20; 3 μM Cy5-dT20, 3 μM unlabeled dT20; each in 50 mM HEPES [pH 7.5], 0.1% βME) and N (4 μM N in 50 mM HEPES [pH 7.5], 20 mM NaCl, 0.1% βME) were used for stocks. In each of these experiments, these stocks were diluted in their background buffers and combined in a 1:1 volumetric ratio to yield the concentrations indicated in the figure.

### Turbidity assays

For the two-dimensional version of the experiment depicted in Figure 2B, a 2x stock solution of 32 μM N in 50 mM HEPES [pH 7.5], 20 mM NaCl, 0.1% βME was combined with 2x solutions of dT20 in 50 mM HEPES [pH 7.5], 0.1% βME in equal volumes. The turbidity of each sample was measured after 15 minutes in a 1 mm pathlength cuvette.

For the phase diagrams in Figure 3A, samples were prepared in clear 384-well microplates (Greiner Bio-One; ref. 781201) by a BioMek FX-P liquid handling robot (Beckman Coulter) at 23^°C^. Stock solutions of 24 and 32 μM N in 50 mM HEPES [pH 7.5], 20 mM NaCl, 0.1% βME were pipetted onto a microplate and serially diluted. A separate stock of 96 μM unlabeled dT20 in 50 mM HEPES [pH 7.5], 0.1% βME was used to prepare a range of dT20 concentrations. N and dT20 were combined in a 1:1 volumetric ratio to yield the concentrations indicated in the figure. To monitor turbidity, we measured the absorbance at 600 nm using a Synergy H1 (BioTek) plate reader over two hours. Assays were performed in triplicate on separate days for statistical analysis.

### Visualization of N Liquid-Liquid Phase Separation by Differential Interference Contrast and Epifluorescence Microscopy

A stock solution of 32 μM N in 50 mM HEPES [pH 7.5], 20 mM NaCl, 0.1% βME was mixed at a 1:1 volumetric ratio with different concentrations of Cy3-dT20 in 50 mM HEPES [pH 7.5], 0.1% βME in a microfuge tube at 23^°C^. Micrographs in each figure were representative of a triplicate dataset.

To visualize liquid-liquid phase separation, 4 μL drop of each sample was pipetted onto a glass slide (VWR) and covered with #1 12 mm diameter cover slip (Premium Line). Samples were imaged using differential interference contrast and Cy3 epifluorescence on a Nikon Eclipse NiE operating with an 80x oil immersion objective. In time-based experiments, fresh slides were prepared at each timepoint.

For condensate disruption experiments, a solution containing 16 μM N, 16 μM Cy3-dT20 was prepared as described previously. After incubation, this solution was diluted with background buffer (50 mM HEPES [pH 7.5], 10 mM NaCl, 0.1% βME), 16 μM Cy3-dT20, or 3.2 μM Cy3-dT20.

### Crosslinking

A 2x stock of 32 μM N in 50 mM HEPES [pH 7.5], 20 mM NaCl was combined in equal volumes with 50 mM HEPES [pH 7.5], a 2x stock of dT20 in the same buffer, or with 50 mM HEPES, 100 mM Tris [pH 7.5] to test for spurious crosslinks. Samples were incubated for 15 minutes before crosslinking with EDC (1-ethyl-3-(3-dimethylaminopropyl)carbodiimide hydrochloride; Pierce) and sulfoNHS (sulfo-N-hydroxylsuccinamide; Sigma Aldrich) or with BS3 (bis(sulfosuccinimidyl)suberate; ThermoFisher). 5x stocks of crosslinkers were prepared in water before adding to samples. EDC/sNHS crosslinking was performed for 2 hours at 23^°C^. BS3 crosslinking was performed for 30 minutes at 23^°C^. Crosslinkers were quenched by adding 150 mM glycine (pH 7.5) for 15 minutes. Samples were boiled for 10 minutes in loading dye containing 10% βME and run on 4-15% Protean SDS-PAGE gels (BioRad). Gels were stained with Coomassie Brilliant Blue R250 and imaged on a BioRad Chemidoc.

## Acknowledgements

We thank Andras Kun of the Light Microscopy Imaging Center and the staff of the Electron Microscopy Center and Physical Biochemistry Instrumentation Facility at Indiana University Bloomington.

