## Supplementary Figures for "A narrow ratio of nucleic acid to SARS-CoV-2 N-protein enables phase separation"

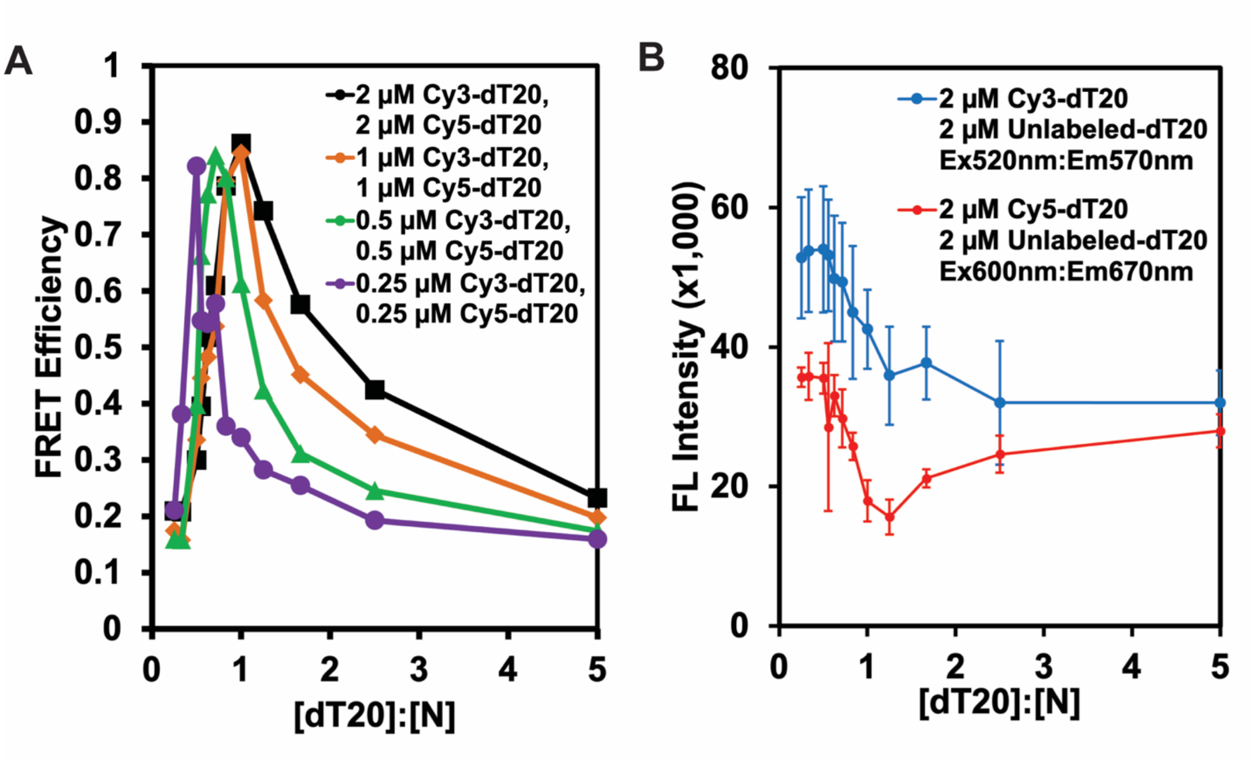


**Supplemental Figure 1. Fluorescently labeled dT20s reveal formation of multiprotein complexes.** (A) FRET efficiency (excitation at 520nm) of N-dT20 complexes using equal parts Cy3- and Cy5-dT20 with different ratios of N. Note that the ratio that produces maximum FRET efficiency depends on the concentration of dT20. (C) Self-FRET and protein-induced fluorescence enhancement for Cy3-dT20 (blue line; excitation at 520 nm, emission at 570 nm) and Cy5-dT20 (red line; excitation at 600 nm, emission at 670 nm) was assessed at a range of [N]:[dT20] ratios

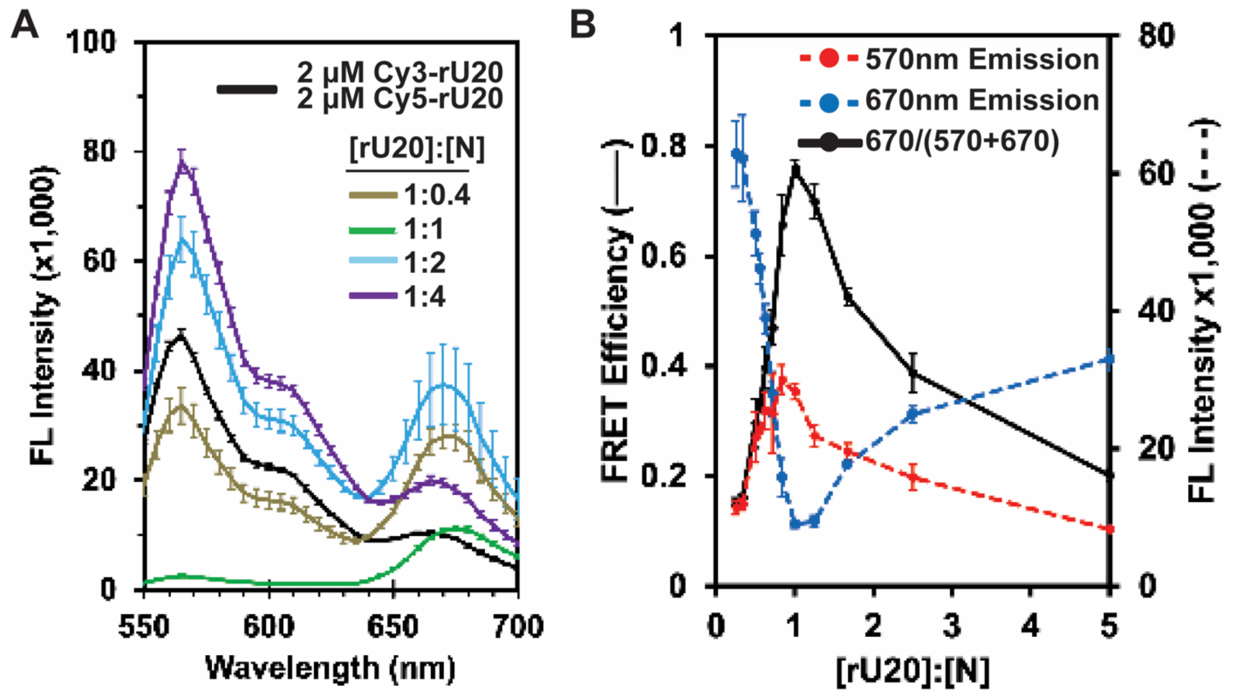


**Supplemental Figure 2.** (B) Emission spectra (excitation at 520nm) of labeled dT20 (2 μM each of Cy3- and Cy5-dT20) with different concentrations of N. Values in the legend represent the molar ratio the total dT20:N. (B) The emission from Cy3-rU20 (blue line) and Cy5-rU20 (red line) shows a minimum near a molar ratio of 1 rU20 to 1 N. This behavior is identical to dT20.

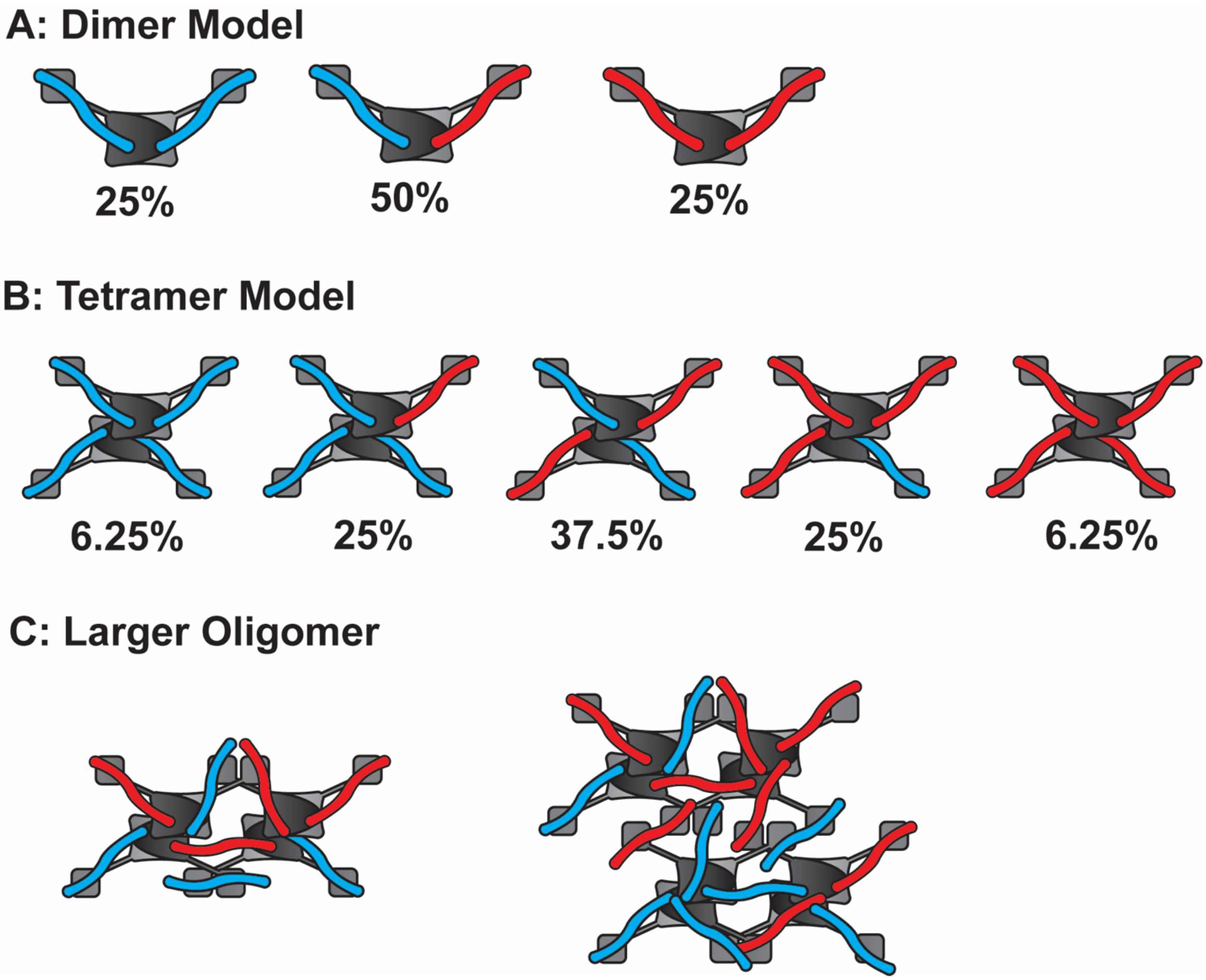


**Supplemental Figure 3.** (A) Illustration of a model in which N forms only a dimer with no further multimerization. Cy3-dT20 and Cy5-dT20 are shown as blue and red sticks, respectively. (B) Model of an N tetramer with four bound dT20s. (C) Model of larger N oligomers. The complexes shown in (B) and (C) have a high likelihood of incorporating both a donor and acceptor fluorophore.

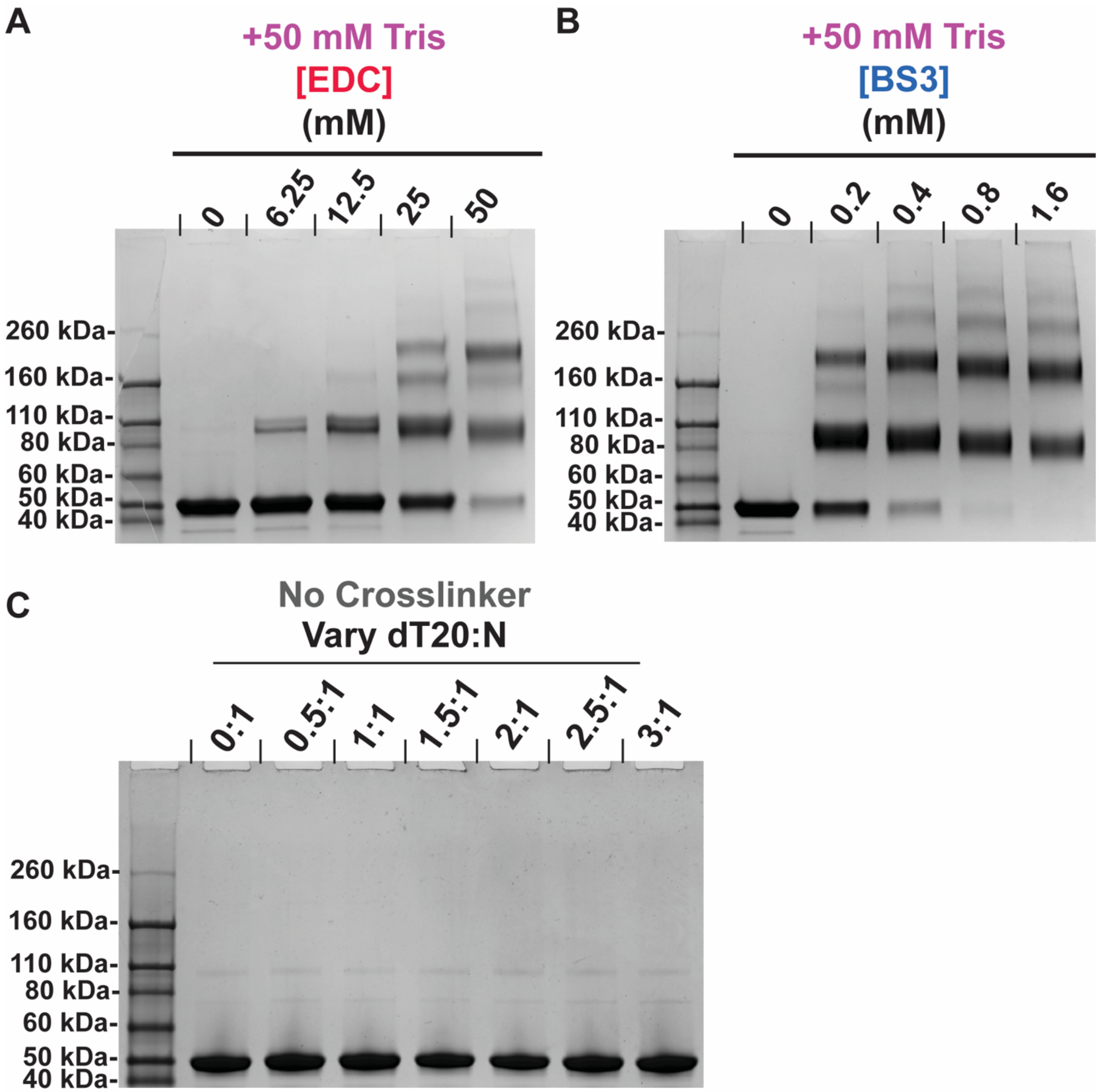


**Supplemental Figure 4.** Crosslinking in the presence of 50 mM Tris for (A) EDC and (B) BS3 shows a similar banding pattern to crosslinking without competitive inhibitor. Bands corresponding to monomer, dimer, tetramer, and higher-MW bands via SDS-PAGE. For EDC crosslinking, sulfoNHS was added to 0.2x the concentration of EDC. (C) SDS-PAGE of N without crosslinker shows no difference in banding pattern with or without dT20.
